# The Ocean Gene Atlas: exploring the biogeography of plankton genes online

**DOI:** 10.1101/271882

**Authors:** Emilie Villar, Thomas Vannier, Caroline Vernette, Magali Lescot, Aurélien Alexandre, Paul Bachelerie, Thomas Rosnet, Eric Pelletier, Shinichi Sunagawa, Pascal Hingamp

**Affiliations:** Sorbonne Universités, UPMC Université Paris 06, CNRS, Laboratoire Adaptation et Diversité en Milieu Marin UMR7144, Station Biologique de Roscoff, Roscoff, France; Aix Marseille Univ, Université de Toulon, CNRS, IRD, MIO UM 110, 13288, Marseille, France; Génomique Métabolique, Genoscope, Institut de Biologie François-Jacob, CEA, CNRS, Univ Evry, Univ Paris-Saclay, 91057 Evry, France; Department of Biology, Institute of Microbiology, ETH Zurich, Zurich, Switzerland

## Abstract

The Ocean Gene Atlas is a web service to explore the biogeography of genes from marine planktonic organisms. It allows users to query protein or nucleotide sequences against global ocean reference gene catalogs. With just one click, the abundance and location of target sequences are visualized on world maps as well as their taxonomic distribution. Interactive results panels allow for adjusting cutoffs for homology and displaying the abundances of genes in the context of environmental features (temperature, nutrients, etc.) measured at the time of sampling. The ease of use enables non-bioinformaticians to explore quantitative and contextualized information on genes of interest in the global ocean ecosystem. Currently the Ocean Gene Atlas is deployed with i) the Ocean Microbial Reference Gene Catalog (OM-RGC) comprising 40 million non-redundant mostly prokaryotic gene sequences, and ii) the Marine Atlas of *Tara* Ocean Unigenes (MATOU) composed of >116 million eukaryote unigenes, both produced by the *Tara* Oceans sequencing effort. Additional datasets will be added upon availability of further marine environmental datasets that provide the required complement of sequence assemblies, raw reads and contextual environmental parameters. Ocean Gene Atlas is a freely-available web service at: http://tara-oceans.mio.osupytheas.fr/ocean-gene-atlas/.

## INTRODUCTION

Marine plankton provide essential ecosystemic functions on the planet: at the basis of the ocean food web, they contribute about half of global primary production (1). Plankton are also key players of biogeochemical cycles in the ocean and drive the biological carbon pump (2). Due to their large distribution (oceans represents 71% of the Earth surface) and their substantial biogeochemical roles, plankton are considered to be main actors in the global climate regulation (3) and also constitute a potential source of innovations for blue biotechnology (4). Because of the difficulties associated with sampling plankton in the open and deep ocean, and because of the ultra high throughput required to sequence the genetic makeup of such complex communities, environmental genomics resources for these elusive organisms have only become available recently (reviewed by Mineta and Gojobori (5)). The first large scale sequencing of marine microbiomes led by the Global Ocean Sampling expedition (6) produced a 6.1 million gene catalog mostly from sunlit ocean prokaryotes.

More recently, the *Tara* Oceans pan-oceanic expedition deployed a holistic sampling of plankton ranging in size from viruses to fish larvae, coupled with comprehensive *in situ* biogeochemical measurements which provide the detailed environmental contexts necessary for ecological interpretation of marine microbiomes (7). Two complementary genesets have been released so far from the *Tara* Oceans sequencing effort: i) The Ocean Microbial Reference Gene Catalog (OM-RGC) and ii) The Marine Atlas of *Tara* Oceans Unigenes (MATOU). The OM-RGC is a comprehensive collection of 40 million genes from viruses, prokaryotes and picoeukaryotes with a size up to 3 μm (8) retrieved from public marine plankton metagenomes and reference genomes. The MATOU is a catalog of 116 million unigenes obtained from poly-A+ cDNA sequencing of different organismal size fractions ranging from 0.8 to 2000 μm (9). About half of the unigenes have a predicted taxonomic assignation representing genes from more than 8000 mostly eukaryotic organisms.

Holistic environmental genomic datasets greatly empower ecological and evolutionary studies that target specific marker or candidate genes known or hypothesized to play a role in biotic interactions or biogeochemical processes (reviewed by Lee et al.(10)). Testing such hypotheses requires detailed analyses of voluminous genomic data in their precise environmental context, a task which requires extensive expertise in interrogating heterogeneous data types (i.e. sequencing reads, assemblies, genes together with their taxonomic and functional annotations, environmental variables) to provide integrated interpretations. In order to allow biologists to easily mine such large and complex datasets without the requirement for either significant hardware or programming skills, we make freely available a web service to visualize the geolocalized abundances of taxonomically annotated plankton genes in the context of environmental features.

## INTERFACE AND FUNCTIONALITY

The Ocean Gene Atlas (OGA) web service provides a submission interface to collect a nucleic or protein sequence query, it then executes data mining procedures on dedicated high performance hardware, and returns interactive result panels for data exploration (Figure 1). A user manual and two case study example sequences are provided online.

**Figure 1.**
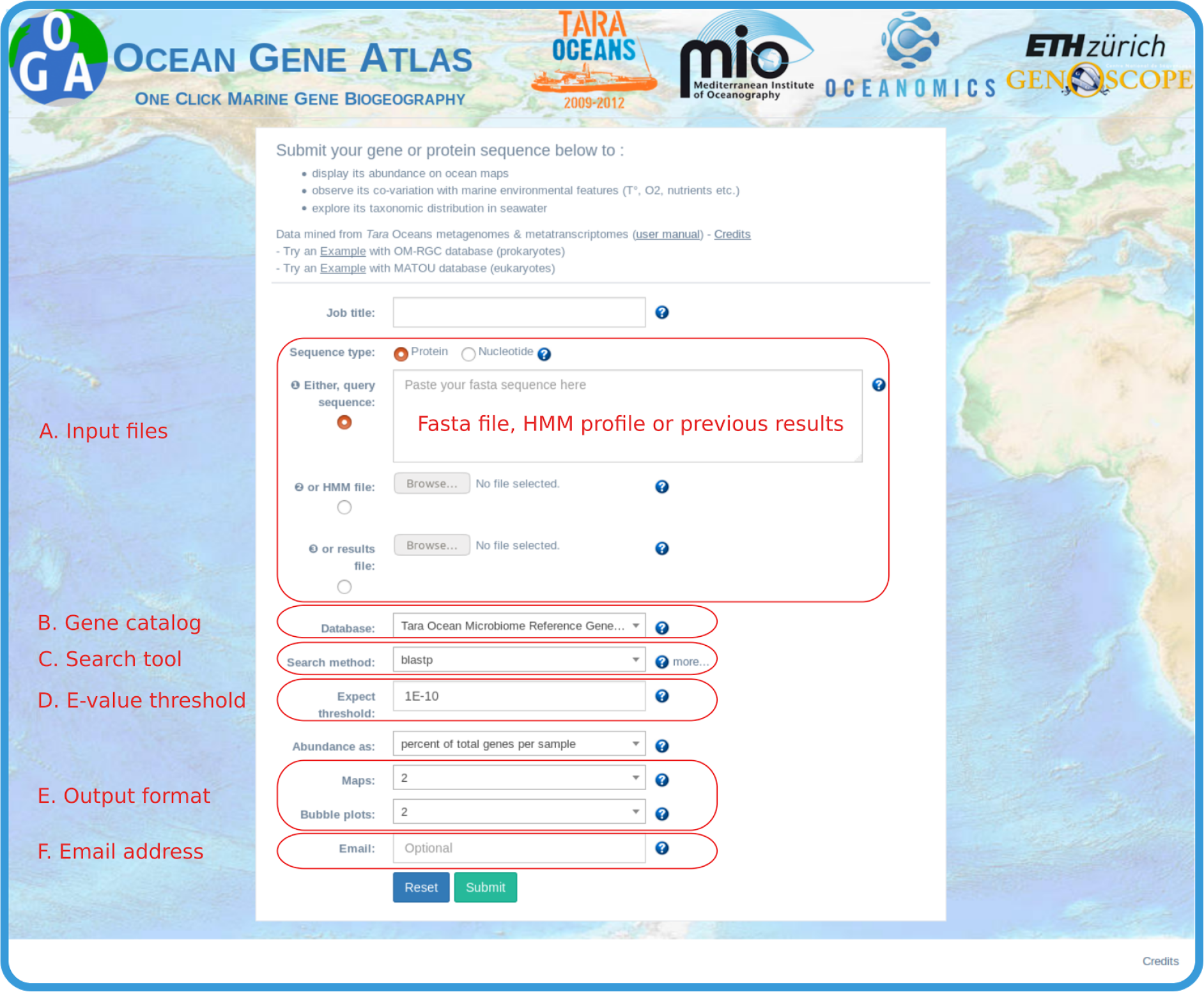
The Ocean Gene Atlas query submission interface (**A**) The query can be either i) a fasta format sequence, ii) an uploaded HMM profile or iii) an uploaded results file from a previous search. (**B**) Two gene catalogs are currently available: OM-RGC, a catalog of mostly prokaryotic genes from plankton metagenomes, and MATOU, a catalog of mostly eukaryotic transcripts from plankton metatranscriptomes. (**C**) The sequence similarity search algorithm is one of Blast, Diamond or hmmer. (**D**) E-value threshold to filter the results. (**E**) Selection of the number of interactive panels in the results page. (**F**) Optionally notification of results availability can be sent by email.

### The query submission interface

The user-provided query (Figure 1A) may be a nucleotide or amino acid sequence in FASTA format, or a hidden Markov model profile (HMM, (11)). Users can search for homologs in either one of two marine gene catalogues (Figure 1B): the OM-RGC (8) or the MATOU (9). Users can select different methods to identify homologs (Figure 1C) depending on the type of query (nucleotide/amino acid sequence or HMM) and the user prefered trade-off between accuracy and speed of the search (12). For protein sequence queries, similarity search tools are blastp (13) and Diamond (14). For nucleotide sequence queries, users can choose to carry out the similarity search against a nucleotide database (blastn), or to translate the nucleotide query to search against a protein database (blastx, diamond blastx). For more sensitive homolog identification, users can provide an HMM instead of a FASTA sequence (either pre-built, e.g. by Pfam (15), or custom-built from protein alignments using the hmmer package, http://hmmer.org). For both sequence similarity and HMM searches, the e-value threshold may be customized (Figure 1D), as well as the number of interactive panels in the results page (Figure 1E). The email field (Figure 1F) is strictly optional since a bookmarkable URL for the results is immediately provided to the user upon submission. Results are usually returned to the user within 30 seconds, but for slower queries (e.g. HMM searches against the larger MATOU catalog which may take up to several minutes or longer in cases of high affluence), the user can choose to provide an email address in order to be notified when results are available (Figure 1F).

### The interactive result panels

The quantitative distribution of environmental sequences presenting similarities to the user query are displayed in three interactive panels: geographic distribution (Figure 2A), co-variation with environmental features (Figure 2B), and taxonomic distribution (Figure 2C). Furthermore, the underlying data necessary to build the figures can be downloaded as tab delimited flat files for further analysis outside of OGA (Figure 2D): list of similarity search hits and corresponding FASTA formatted sequences, gene per sample abundance matrix, as well as contextual environmental features for each sample. The set of similarity search hits that are included in the three interactive panels (referred hereafter as homologs) can be interactively filtered by click-and-drag adjustment of the E-value threshold directly over the provided E-value distribution histogram. The world maps of Figure 2A display quantitative geographical distributions of the homologs as filled circles with sizes proportional to their combined abundance at the user-selected sampling depth, whilst circle colors indicate the size fractionation applied to the sample (e.g. [0.2-3μm] represents plankton collected on 0.2μm pore membranes after a 3μm prefiltration step). The meaning of the acronyms and references to source databases are provided in a user guide hyperlinked on the results page. The side-to-side display of multiple maps enables abundances comparisons between distinct size fractions and/or sampling depths. Co-variation of gene abundances and environmental features can be examined on bubble plots (Figure 2C) for user selectable combinations of sampling depth and size fractions. Finally, the taxonomic distribution of the target genes are displayed in multi-layered and interactive Krona pie-charts (16) either for each distinct sample (by clicking on the circles in the world maps) or for the full dataset (Figure 2C). The charts displayed on the Ocean Gene Atlas results page can be annotated online and downloaded as image files in vector graphics formats (SVG and PDF) suitable for publication.

**Figure 2.**
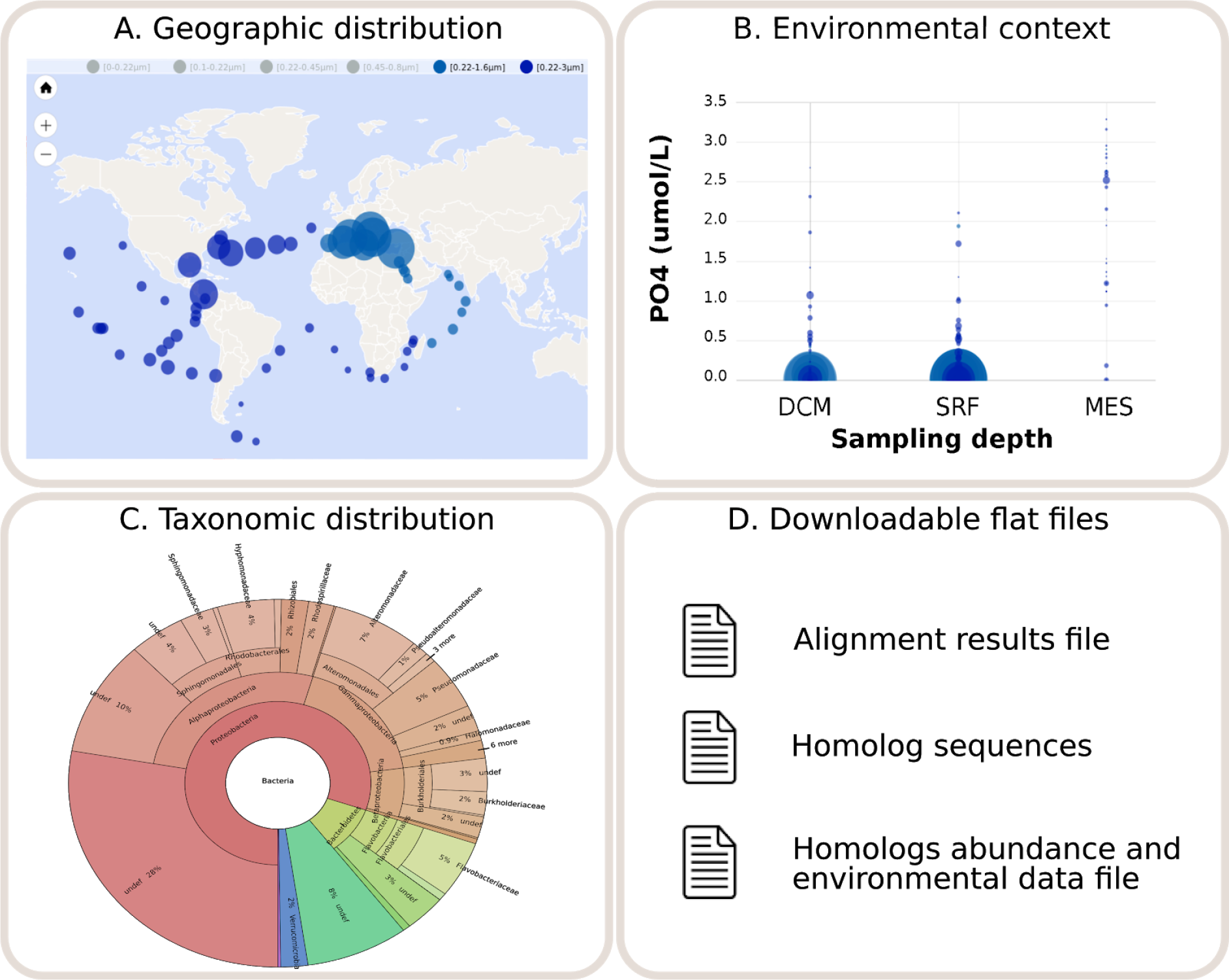
The Ocean Gene Atlas interactive results panels. (A) Homolog abundances are represented by the diameter of filled circle for each sample at user selected sampling depths (e.g. subsurface or mesopelagic). Circle colors represent the organismal size fractions (e.g. [0.2-3μm]). (B) Co-variation of homolog abundances with specific environmental variables are shown on bubble plots for each sampling depths: subsurface (SRF), deep chlorophyll maximum (DCM) and mesopelagic (MES). (C) Taxonomic distribution of the homolog genes’s predicted origins are represented on interactive Krona plots. (D) Result files can be downloaded as tab delimited flat files.

## DATA SOURCES

Datasets suitable for inclusion in the Ocean Gene Atlas require three complementary data objects: gene sequence catalogs, gene abundances in samples, and sample environmental context (Figure 3).

**Figure 3.**
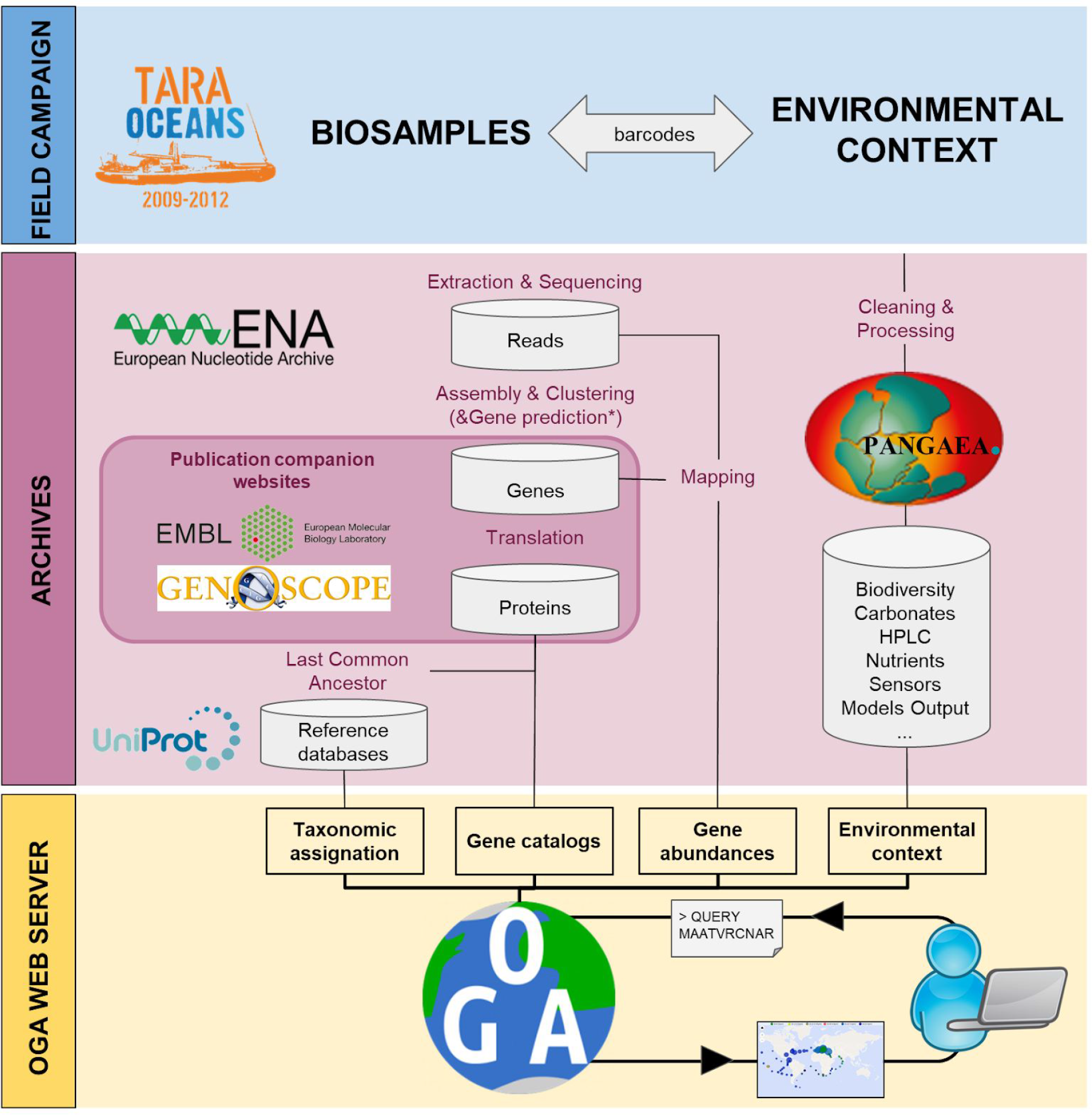
Data sources for the the Ocean Gene Atlas workflow. Field campaigns (blue) have collected plankton biosamples and measured *in situ* environmental parameters. The OGA web server (yellow) combines heterogeneous data published by distinct archives (pink): EBI ENA for sequencing reads, published articles companion websites for gene catalogs and taxonomic annotations, PANGAEA for contextual environmental data.

### Gene catalogs

The building of the OM-RGC and MATOU gene catalogs included in the first release of OGA are detailed in their corresponding release articles (8, 9 respectively). Briefly, to construct OM-RGC, 7.2 terabases of plankton metagenome shotgun sequencing reads were assembled for 243 *Tara* Oceans samples. Genes were predicted in the assemblies and were clustered at 80% sequence identity together with genes from publicly available marine genomic and metagenomic datasets to generate a non-redundant set of 40 million reference genes. These genes were translated and taxonomically annotated by retrieving the last common ancestor of homologs identified in reference protein sequence databases. The MATOU catalog was obtained from the assembly of 16.5 terabases of plankton metatranscriptome (cDNA sequences corresponding to polyA+ enriched RNA), representing 441 *Tara* Oceans samples. The subsequent contigs obtained for each assembled sample were then clustered at 95% sequence identity to construct a catalog of 116.8 million transcribed sequences. Due to the difficulty of accurate eukaryotic gene calling from low coverage metatranscriptomes, the proteic version of the MATOU catalog was obtained by six frame translation of the nucleic MATOU gene catalog using the sixpack package from the EMBOSS suite (17). About half of the unigenes were taxonomically annotated using a similar last common ancestor approach as described above for the OM-RGC.

### Gene abundances

The abundance of each catalog gene in specific biosamples was estimated by evaluating the coverage of raw sequencing reads mapped to the gene’s nucleotide sequence. *Tara* Oceans sequencing reads from 243 metagenomes corresponding to the smallest size fractions (0-0.22 μm, 0. 1–0.22 μm, 0.22–0.45 μm, 0.45–0.8 μm, 0.22–1.6 μm and 0.22–3 μm) were mapped onto OM-RGC, whilst the reads from 441 metatranscriptomes corresponding to the largest size fractions (0.8–5 μm, 5–20 μm, 20–180 μm and 180–2000 μm) were mapped onto MATOU. When OM-RGC is queried, abundance estimates may be expressed in one of two available normalization schemes: i/ the gene’s read coverage is divided by the sum of the total gene coverages for the sample (“*percent of total genes per sample*“), or ii/ the gene’s read coverage is divided by the median of the coverages of a set of ten universal single copy marker genes (“*average copies per cell*”) that were previously benchmarked for their suitability for metagenomics data analysis (18). MATOU gene abundance estimates are expressed as gene read coverage computed in RPKM (Reads Per Kilobase covered per Million of mapped reads) divided by the sum of the total gene coverages for the sample (“*percent of total genes per sample*“).

### Environmental context

Contextual environmental parameters are linked to the sequence datasets via barcodes assigned to each *Tara* Oceans sample (19). These metadata serve to geo-localize the samples and provide biogeochemical characteristics of the sampled seawater. Environmental parameters used by the OGA web service are obtained from PANGAEA (https://doi.org/10.1594/PANGAEA.875582), the open access library which archives and distributes georeferenced data from earth system research. The environmental parameters provided by OGA (Figure 3) are either classical oceanographic measures obtained *in situ* (e.g. depth, salinity, temperature, oxygen, chlorophyll a, etc.) or mesoscales features estimated from oceanographic models and remote satellite observations (e.g. nutrient concentration at 5m depth or net primary production). Estimated values are indicated by a star in the drop-down menu of bubble plot panels (Figure 2C). Descriptions of the environmental parameters available in OGA as well as corresponding PANGAEA hyperlinks are provided in the OGA_user_manual hyperlinked from the OGA results page.

## IMPLEMENTATION

The Ocean Gene Atlas web server is implemented through a classical Model-View-Controller pattern architecture using the Laravel 5.4 PHP framework. Developed on the GNU/Linux, the application server communicates with the user through an Apache HTTP server using HTML5, CSS3, JS, BLADE and AJAX to retrieve user requests and display results. The PHP application server queries abundance and environmental data stored in a SQLite3 relational database.

## CASE STUDY

Upon phosphorus deficiency, bacterioplankton have established a widespread strategy of replacing membrane phospholipids with alternative non-phosphorus lipids. Sebastián *et al.* (20) have shown that this response is conserved among diverse marine heterotrophic bacteria. Several experiments of mutagenesis and complementation have then confirmed the roles of the phospholipase C (PlcP) and a glycosyltransferase in lipid modelling. Analyses of metagenome datasets such as the Global Ocean Sampling (GOS) and *Tara* Oceans have confirmed that PlcP is abundant in low phosphate concentrations areas. We reproduced the analysis of Sebastián et al. (20) using OGA to verify that the web service conforms to the published results. We used the same phospholipase C (EAQ46983) as a BLASTp query sequence with the author’s e-value threshold of 1e^−40^ to search for homologous sequences in the OM-RGC catalog (the same metagenome dataset used by Sebastián *et al.*(20)). The 952 PlcP homologs identified showed higher abundances in Mediterranean subsurface samples (Figure 4A) related to low phosphorus concentration (Figure 4B) and mostly originated from Proteobacteria and Bacteroidetes (Figure 4C), which agrees with the previously published interpretations of Sebastián *et al.* that marine heterotrophic bacteria display reduced phosphorus requirements upon phosphorus deficiency by PlcP-mediated replacement of membrane phospholipids by alternative non-phosphorus lipids.

**Figure 4.**
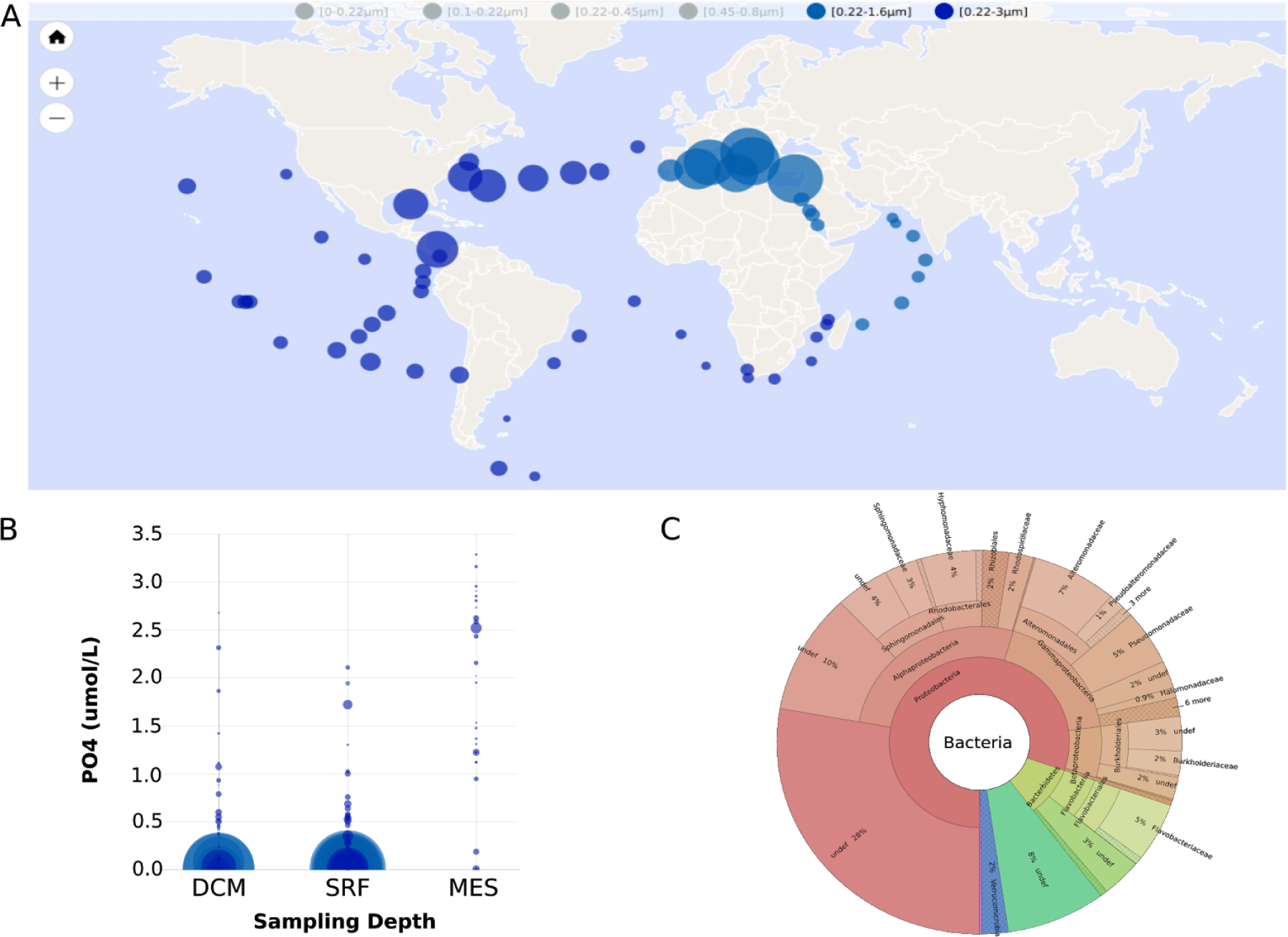
Phospholipase C (PlcP) biogeography produced by the Ocean Gene Atlas web service (A) Abundance of PlcP homologs in the OM-RGC subsurface samples. (B) Bubble plot of the PlcP abundance in relation to PO4 concentrations; DCM: Deep Chlorophyll Maximum layer, SRF: subsurface and MES: mesopelagic zone. (C) Krona plot of the taxonomic distribution of the PlcP homologs.

## CONCLUSION/PERSPECTIVES

By hiding the complex and time consuming integration of heterogeneous data sources behind a user-friendly minimalist web form, the Ocean Gene Atlas web server has the potential to broaden the access to the rapidly accumulating environmental marine genomics datasets. Enabling marine biologists to mine such data - without specific high performance hardware or programming skills - is one of the keys to extract knowledge and understanding from these valuable but underexploited resources. With the current first release of OGA, users can search protein or nucleotide sequences homologs in two of the largest marine gene catalogs representing all three eukaryotic, prokaryotic and viral domains. These two first catalogs will be periodically updated as further marine environmental genomics databases are publically released. The prerequisites for inclusion in OGA are the availability of the three core resources: gene sequence catalogs, gene abundance estimates in biosamples, and geolocalized environmental context of biosamples. Leveraging the total 2100 *Tara* Oceans biosamples sequenced so far (21), our short term roadmap is to i) extend the *Tara* Oceans datasets by including further sampling sites from the Polar Circle expedition, ii) complement the eukaryotic MATOU metatranscriptome catalog with corresponding metagenomic abundances, and iii) complement the OM-RGC with prokaryotic metatranscriptomes.

## AVAILABILITY

Ocean Gene Atlas is a freely-available web service at: http://tara-oceans.mio.osupytheas.fr/ocean-gene-atlas/.

## ACCESSION NUMBERS

*Tara* Oceans shotgun sequencing have been deposited with the European Nucleotide Archive at EBI under accession number PRJEB402 (OM-RGC) and PRJEB6609 (MATOU). The predicted genes from the OM-RGC have been deposited with the European Nucleotide Archive at EBI under the accession numbers ERZ096909 to ERZ097151, and the protein sequences are available at: ftp://ftp.genome.jp/pub/db/mgenes/Environmental/Tara.pep.gz. All MATOU resources are available at http://www.genoscope.cns.fr/tara/. Registry of all the samples from the *Tara* Oceans Expedition (2009-2013) have been deposited at *PANGAEA*: https://doi.org/10.1594/PANGAEA.875582.

## ACKNOWLEDGEMENT

This article is contribution number XX of the *Tara* Oceans expedition.

